# hoodscanR: profiling single-cell neighborhoods in spatial transcriptomics data

**DOI:** 10.1101/2024.03.26.586902

**Authors:** Ning Liu, Jarryd Martin, Dharmesh D Bhuva, Jinjin Chen, Mengbo Li, Samuel C. Lee, Malvika Kharbanda, Jinming Cheng, Ahmed Mohamed, Arutha Kulasinghe, Yunshun Chen, Chin Wee Tan, Fuyi Li, Jose M Polo, Melissa J Davis

**Affiliations:** South Australian immunoGENomics Cancer Institute (SAiGENCI), Faculty of Health and Medical Sciences, The University of Adelaide, Adelaide, SA 5005, Australia; Bioinformatics Division, The Walter and Eliza Hall Institute of Medical Research, Parkville, Melbourne, Victoria 3052, Australia; Department of Medical Biology, Faculty of Medicine, Dentistry and Health Sciences, University of Melbourne, Parkville, VIC 3010, Australia; Adelaide Centre for Epigenetics, School of Biomedicine, Faculty of Health and Medical Sciences, The University of Adelaide, SA 5005, Australia; Frazer Institute, Faculty of Medicine, The University of Queensland, Brisbane, Queensland 4102, Australia; Department of Anatomy and Developmental Biology, Monash University, Wellington Road, Clayton, VIC 3800, Australia; Computational Biology, Isomorphic Labs, Bishopsgate, London, England, EC2M 4RB, United Kingdom

## Abstract

Understanding complex cellular niches and neighborhoods have provided new insights into tissue biology. Thus, accurate neighborhood identification is crucial, yet existing methodologies often struggle to detect informative neighborhoods and generate cell-specific neighborhood profiles. To address these limitations, we developed hoodscanR, a Bioconductor package designed for neighborhood identification and downstream analyses using spatial data. Applying hoodscanR to breast and lung cancer datasets, we showcase its efficacy in conducting detailed neighborhood analyses and identify subtle transcriptional changes in tumor cells from different neighborhoods. Such analyses can help researchers gain valuable insights into disease mechanisms and potential therapeutic targets.

## Introduction

Spatial transcriptomics stands out as a powerful technology, offering a distinctive perspective that goes beyond traditional bulk RNA-seq and single-cell RNA-seq (scRNA-seq) methods. Since it conserves the spatial information of a tissue, it yields valuable insights into the complex molecular and cellular landscapes, uncovering spatial variations and relationships often overlooked by conventional approaches. Recent advancements in spatial-omics platforms, including Nanostring CosMx Spatial Molecular Imager (1), 10X Genomics Xenium (2), Vizgen Merscope (3), Akoya CODEX (4,5), and others, have facilitated the generation of single-cell level spatial data.

However, despite the potential of spatial transcriptomics, the field is still in its early stages, with many analyses resembling conventional scRNA-seq-like approaches. These analyses often disregard the rich spatial context of the data, failing to harness the cellular coordinates. Thus, this shift towards high-resolution spatial profiling and the lack of appropriate methods has created a pressing demand for innovative analytical tools capable of fully exploiting these datasets. Cellular neighborhood analysis, a powerful approach to fully utilize cell spatial information, becomes particularly important when applied to single-cell level spatial transcriptomics data. Bioinformatics tools are needed to identify and characterize the niches or neighborhoods in which cells reside, as these regions may harbor crucial tissue micro-environment (TME) biology that influences the fundamental tissue biology, physiology as well as responses to therapy and disease progression. Therefore, understanding these neighborhoods is key for the full utilization of the spatial data and to provide researchers with novel insights into cellular interactions and communications within the TME, offering a nuanced understanding of the complex biological processes at play. Such insights hold the potential to enhance our understanding of complex diseases like cancer and contribute to the development of more effective therapies.

In recent years, there has been a growing trend in the development of methods dedicated to conducting neighborhood analyses to interpret complex cellular neighborhoods within spatial transcriptomics data (Table 1). These methods range from clustering-based approaches that leverage frequency matrices of k-nearest cells (1) to graph network-based strategy that is built into interactive viewer (6). Widely used toolkits Squidpy (7) and Giotto (8) have made substantial contributions to the field by facilitating neighborhood analysis via enrichment tests using a graph-based approaches compatible across multiple spatial technology platforms. Additionally, many tools have been developed to detect spatial domains from spatial transcriptomics datasets by accounting for the spatial information, i.e. cellular neighborhood when clustering data via various approaches, including BuildNicheAssay from Seurat (9), MERINGUE (10), BANKSY (11), BayesSpace (12), STAGATE (13), SpaGCN (14) and UTAG (15).

**Table 1.** Features of existing neighborhood/domain identification methods for spatial transcriptomics data.

| Features | Giotto(8) | Squidpy(7) | Seurat | Banksy | BayesSpace | MERINGUE | SpaGCN | Stagate | Utag | hoodscanR |
| --- | --- | --- | --- | --- | --- | --- | --- | --- | --- | --- |
| Language | R | Python | R | R/Python | R | R | Python | Python | Python | R |
| Infrastructure | Giotto object | AnnData | Seurat object | Spatial Experiment | Spatial Experiment | NA | AnnData | AnnData | AnnData | Spatial Experiment |
| Multi-platforms compatible | Yes | Yes | Yes | Yes | Yes | Yes | Yes | Yes | Yes | Yes |
| Co-localization | Yes | Yes | No | No | No | No | No | No | No | Yes |
| Multi-neighborhood membership | No | No | No | No | No | No | No | No | No | Yes |
| Cell-level neighborhood profiles | No | No | No | No | No | No | No | No | No | Yes |
| Spatial domain detection | Yes | No | Yes | Yes | Yes | Yes | Yes | Yes | Yes | Yes |

Nevertheless, despite these advancements, there are critical gaps in existing methodologies. Most notably, while some existing tools can detect spatial domains that comprise multiple cell types, such as UTAG, SpaGCN and Giotto’s HMRF-based approach, they do not provide partial membership at a single-cell level. For example, when cells reside in neighborhoods characterized by a mixture of B cells and stromal cells, current methods tend to categorize such neighborhoods as either exclusively B cell or stromal cell neighborhoods, failing to capture the nuanced composition of cellular environments. Furthermore, current tools lack the capability to provide cell-level neighborhood annotations, meaning detailed neighborhood profiles for individual cells are unavailable. This critical feature is essential for a comprehensive characterization of the spatial context surrounding each cell. In response to these unaddressed challenges, we developed hoodscanR, a Bioconductor R package designed to perform comprehensive neighborhood analyses on spatial transcriptomics data. Unlike existing methods, hoodscanR aims to bridge critical gaps by enabling per-cell partial membership across multiple neighborhoods, providing a more precise and detailed understanding of the tissue microenvironments. Additionally, hoodscanR generates cell-level neighborhood profiles, a unique feature that allows for an in-depth summarization of the spatial context at a single-cell resolution. Moreover, hoodscanR can identify neighborhood-based spatial domains, offering insights into the higher-order organisation of tissues. In this study, we introduce the functionalities and capabilities of hoodscanR and demonstrate its utility in investigating the cellular neighborhoods within publicly available spatial transcriptomics datasets.

## Materials and methods

### Data pre-processing

Both CosMx and Xenium datasets underwent a rigorous quality control process to ensure the inclusion of high-quality cells in the neighborhood analysis. For the CosMx data, thresholds were set at the 0.1 quantile to filter out cells with low transcript count or low gene detection count across all cells per tissue slide. Additionally, genes with mean expression (log-scaled count per million) and variance lower than the negative probes were excluded from further analyses. In the case of the Xenium data, filtering followed the guidelines outlined in the Squidpy(7) toolkit tutorial. Cells with a transcript count less than 10 and genes detected in fewer than 5 cells were removed from the neighborhood and downstream analyses. As a result, almost 90,000 cells per slide with 870 genes and 156,224 cells with 313 genes were kept for the NSCLC and breast cancer datasets, respectively.

### Cell type annotation

Cell type annotations for the Nanostring CosMx NSCLC data was carried out with modifications as previosuly described in Tan et al 2024 (16). Briefly, specific modifications include using *SCTransform* from the Seurat package (17) to normalise filtered counts from the quality control step. By modelling negative probe detection as a fixed factor, we regressed out the confounding effects caused by background. The annotation process of the data were performed using InSituType (18), with the Single-cell Lung Cancer Atlas (LuCA)(19) as the reference. In terms of 10X Xenium data, cell type annotation was obtained from the Janesick A*, et al* paper (2).

### Metrics calculation for neighborhood probability distribution

Perplexity serves as a fundamental metrics for summarizing the neighborhood probability distribution generated by hoodscanR. It provides a measure of effective diversity of complexity within a cell’s neighborhood. The perplexity 𝑃(𝑥) for a given cell 𝑥 is calculated as:

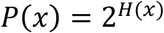

Where 𝐻(𝑥) represents the Shannon entropy (20) of the neighborhood probability distribution of cell 𝑥, defined as:

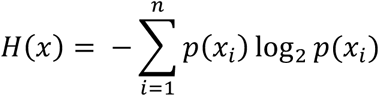

Where 𝑝(𝑥_𝑖_) is the probability of cell 𝑥 located in the 𝑖-th neighborhood and 𝑛 is the total number of distinct neighborhoods. Higher perplexity values indicate greater diversity or complexity within the cellular neighborhoods, suggesting that a larger number of distinct cell types are contributing to the neighborhood. In hoodscanR, perplexity can be calculated using the *calcMetrics* function.

To assess the statistical significance of the observed perplexity values within cell neighborhoods, we employed an empirical permutation test. For each neighborhood, we generated a distribution of perplexity values by randomly shuffling the spatial coordinates of cells and recalculating the perplexity across 1,000 permutations. The empirical p-value for each neighborhood was then calculated as the proportion of permuted perplexity values that were greater than or equal to the observed perplexity value, adjusted for the finite number of permutations:

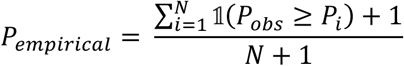

where 𝑃_𝑜𝑏𝑠_ is the observed perplexity for a given neighborhood, 𝑃_𝑖_ represents the perplexity from the 𝑖-th permutation, 𝟙(. ) is an indicator function that equals 1 when the condition inside is true and 0 otherwise, and 𝑁 is the total number of permutations. This correction ensures that the empirical p-values are properly calibrated, even with a limited number of permutations, thus providing a robust measure of statistical significance.

### Hyperparameter k and 𝜏 testing

In order to test the effect different parameter k values have on the results of hoodscanR, we first test a range of k values (10, 50, 100, 200, 500, 1000), using the default 𝜏 setting from the *scanHoods* function. We then computed the Pearson correlation between the resulting probability matrices to assess consistency of the outcomes across different k. For testing the 𝜏 parameter, we fixed k = 100 and examined a range of 𝜏 values, which were derived from different scaling of the distance matrix (see Availability of data and materials).

### Benchmarking of co-localization analysis

Both 10X Xenium breast cancer and Nanostring CosMx datasets were utilized for benchmarking the co-localization analysis among hoodscanR, Squidpy, and Giotto. To ensure robustness, the data were randomly subset into ten different sizes ranging from 0.1 to 1, after which each package’s respective methodologies were applied.

Specifically, for hoodscanR, neighborhood identification and co-localization analysis were conducted using the *plotColocal* function. In contrast, Squidpy and Giotto performed network graph construction using the *gr.spatial_neighbors* and *createSpatialDelaunayNetwork* functions, followed by co-localization analysis using the *gr.nhood_enrichment* and *cellProximityEnrichment* functions, respectively.

### Benchmarking spatial domain identification

To benchmark hoodscanR against other state-of-the-art methods in identifying spatial domains from spatial transcriptomics datasets, we selected 12 publicly available datasets. These included 3 CosMx NSCLC, 6 MERFISH mouse colon, 1 STARmap mouse cortex, and 2 Xenium breast cancer slides. Due to the large size of the Xenium breast cancer datasets (555,579 cells and 353,783 cells, respectively), some tools were unable to process these datasets efficiently. To address this, we downsampled these slides to 10,000 and 50,000 cells, while maintaining the original cell type proportion distributions, resulting in a total of 16 datasets. This was achieved by randomly sampling cells from each cell type cluster based on their proportional weighting (see Availability of data and materials). Annotated region labels or pathological annotations in these datasets were used as the ground truth for spatial domains.

Normalization of the gene expression data was handled differently depending on the dataset. For the MERFISH dataset, we directly used the provided log-normalized counts. For all other datasets, we used the *quickCluster* and *calculateSumFactors* functions from the *scran* R package (21) to estimate size factors, followed by the *logNormCounts* function from the scuttle R package (22) to normalize the counts.

We compared hoodscanR against seven other methods capable of performing spatial domain detection: BuildNicheAssay from Seurat, Banksy, BayesSpace, MERINGUE, SpaGCN, Stagate, and Utag. For each method, we calculated a composite performance score to assess the accuracy of spatial domain identification. This score was computed as the mean of four key metrics: Adjusted Rand Index (ARI), Normalized Mutual Information (NMI), purity, and homogeneity. It is defined as:

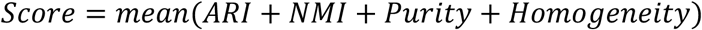

These metrics were chosen for their ability to quantify different aspects of clustering accuracy. ARI measures the similarity between the predicted and true clusters, adjusting for chance. It is defined as:

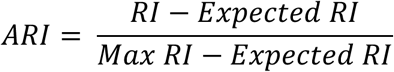

Where RI is the Rand Index, which counts the number of correct pairwise classifications between the predicted and true labels. ARI in this paper is calculated using the *aricode* R package.

NMI quantifies the amount of information shared between the predicted and true clusters. It is defined as:

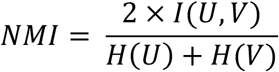

where 𝐼(𝑈, 𝑉) is the mutual information between clusters 𝑈 and 𝑉, and 𝐻(𝑈) and 𝐻(𝑉) are the entropies of the true and predicted clusters, respectively. NMI in this paper is calculated using the *aricode* R package.

Purity measures the extent to which each cluster contains only members of a single class. It is calculated as:

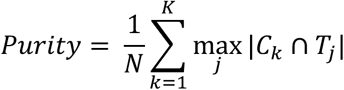

where 𝐶_𝑘_ are the predicted clusters, 𝑇_j_ are the true clusters, and 𝑁 is the total number of samples.

Homogeneity ensures that all the clusters contain only data points which are members of a single class. It is defined as:

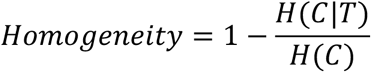

where 𝐻(𝐶|𝑇) is the conditional entropy of the true clusters given the predicted clusters, and 𝐻(𝐶) is the entropy of the predicted clusters.

To ensure fairness in the benchmarking process, all methods were executed using their default settings (see Availability of data and materials). Additionally, a time penalty was applied: if a method failed to complete the processing of a dataset within 24 hours, the process was terminated. This ensures that the comparison accounts not only for accuracy but also for computational efficiency.

### Unsupervised clustering of neighborhood distribution

To perform unsupervised clustering of the neighborhood distribution, the identified neighborhood distribution of each cell was utilized as input data. The K-means clustering algorithm, a widely used method for partitioning data into distinct clusters based on dissimilarity, were used. Specifically, we set the parameters for K-means clustering as iter_max = 1000, nstart = 5, and algo = "Hartigan-Wong". The iter_max parameter determines the maximum number of iterations allowed to converge to a solution, while nstart specifies the number of initial cluster centers to use in the algorithm. Additionally, the "Hartigan-Wong" algorithm was chosen as the method for center initialization. To determine the optimal number of clusters (k) for the K-means clustering algorithm, we used the elbow method. In a nutshell, we first calculated the within-cluster sum of squares for a range of k values and then identified the point at which the distortion or inertia starts decreasing in a linear fashion.

### Differential expression analysis

To conduct the differential expression (DE) analysis on the CosMx NSCLC dataset and identify DE genes among tumor cells from distinct cellular neighborhoods, pseudo-bulk via summation samples were initially generated from cells within the identified neighborhood-based clusters using *summarizeAssayByGroup* function from the *scuttle* R package (22). Subsequently, the *standR* package (23) was used to assess relative log expression (RLE) and perform principal component analysis (PCA) to explore the technical and biological variation in the pseudo-bulk data. Following this, the limma-voom pipeline (24) was utilized for DE analysis with TMM normalisation (25), incorporating slide information as a covariate in the linear model to account for slide-related variations. The resulting statistic was an empirical Bayes moderated t-statistic. Multiple testing adjustment using the Benjamini–Hochberg procedure was then applied to identify DE genes that reached statistical significance (FDR < 0.05). To identify DE genes in the 10X VisiumHD mouse brain dataset, we first extracted neurons from the *Neurod6+* and *Calb2+* neighborhoods and then applied Seurat’s *FindAllMarkers* function (9). We retained only genes meeting two criteria: a log2 fold change greater than 1 and an adjusted p-value less than 0.05. The top ten genes from each group, based on these filters, were subsequently selected for heatmap visualization using the *DoHeatmap* function from Seurat.

### Gene-set enrichment analysis and visualization

Gene-sets from the Molecular Signatures Database (MsigDB, v7.2), including Hallmarks, C2 (curated gene-sets), and C5 (gene ontology terms) categories, along with KEGG pathway gene-sets, were obtained using the *getMisgdb* and *appendKEGG* functions from the msigdb R package (v1.1.5). Gene-set enrichment analysis (GSEA) was performed using *fry* from the limma package (v3.58.1). A false discovery rate of 0.05 was used as the threshold for determining significantly enriched gene-sets. The results of GSEA were systematically examined and visualized using an unbiased approach through the novel network enrichment and visualization R package vissE (26).

## Results

### Development of hoodscanR

hoodscanR uses an efficient computational pipeline to investigate spatial neighborhood relationships among cells within spatial transcriptomics datasets (Figure 1). At the core of hoodscanR, the searching process for nearest cells is initiated using an Approximate Nearest Neighbor (ANN) search algorithm (27), which uses k-dimensional tree to efficiently manage the two-dimensional spatial coordinates of spatial transcriptomics data, providing rapid identification of nearest neighbors while maintaining high accuracy. This facilitates the identification of the k-nearest neighboring cells for each cell in the dataset. This process outputs a list of indices representing the nearest neighbors of each cell, denoted as:

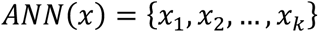

**Figure 1.**
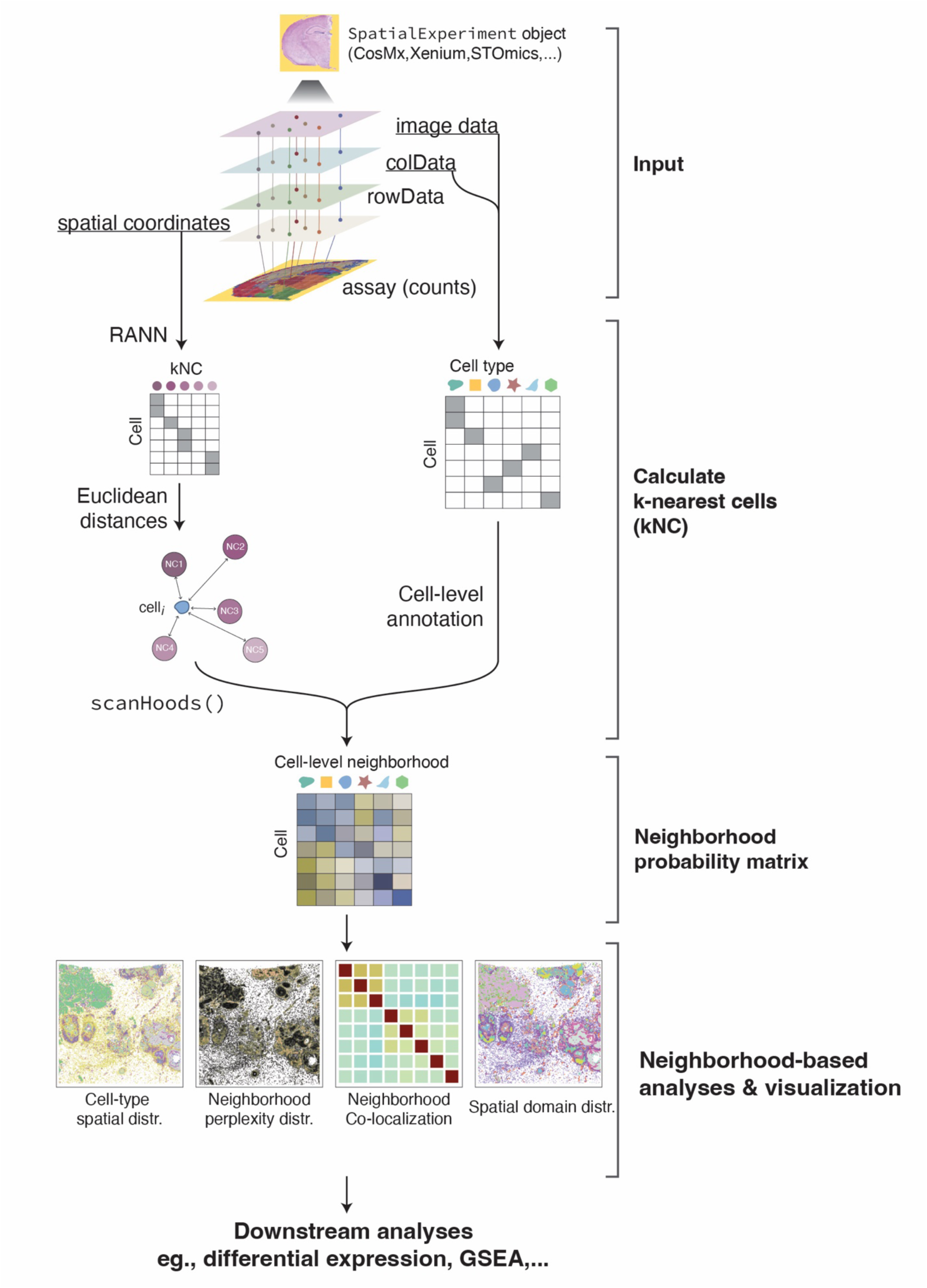
Schematic visualization showing the main components and computational workflow of the hoodscanR package. The process begins with inputting a SpatialExperiment object, which contains spatial data and associated metadata. Next, the package calculates the k-nearest cells based on spatial proximity. This step follows by generating a neighborhood probability matrix, which quantifies the likelihood of cell interactions within their local neighborhood. Finally, the package performs visualizations and downstream neighborhood-based analyses to provide insights into spatial patterns and relationships.

Following the identification of nearest neighbors, hoodscanR calculates the distance between each cell and its k-nearest neighbors. Here Euclidean distance is used due to its simplicity and effectiveness in measuring distances between points in a two-dimensional space. This results in a distance matrix 𝐷, where each element 𝐷_𝑖j_ represents the distance between cell 𝑥_𝑖_ and its neighbor 𝑥_j_ from 𝐴𝑁𝑁(𝑥) (Figure 1). Simultaneously, cell-level annotations provided by users, such as cell types, are used to construct a cell annotation matrix 𝐴, which describes the organisation of cells based on their distances to neighboring cells. Each entry 𝐴 indicates whether cell 𝑥_𝑖_ belongs to annotation group 𝑗:

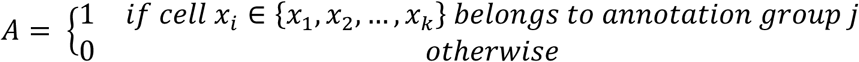

The fundamental function of hoodscanR is to identify cellular neighborhoods within spatial transcriptomics data. It achieves this by using the SoftMax function, enhanced by a hyperparameter 𝜏 (tau), which governs the shape of the resulting probability distribution and provides control over the influence of neighboring cells. The algorithm is expressed as follows:

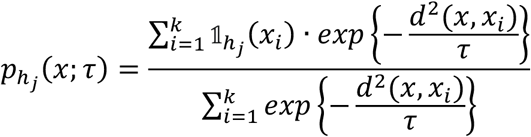

where 𝑥, 𝑥_𝑖_ ∈ {𝑥_1_, 𝑥_2_, … , 𝑥_𝑘_}, and

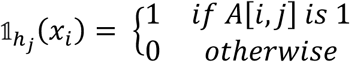

Where:

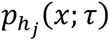 denotes the probability of cell 𝑥 residing within the local neighborhood ℎ_j_.

𝑑(𝑥, 𝑥_𝑖_) signifies the spatial Euclidean distance between cell 𝑥 and its neighboring cell 𝑥_𝑖_.

𝜏 stands as the hyperparameter, facilitating fine-tuned modulation of the impact of neighboring cells.

ℎ_j_ denotes the cell neighborhood 𝑎, defined by the cell-level annotations provided by users. For example, if cell types were provided, ℎ_j_ means cell type 𝑗 neighborhood.

𝟙(. ) is the indicator function, which checks whether cell 𝑥_𝑖_ belongs to the neighborhood ℎ_j_ as per the annotation matrix 𝐴.

Upon the aggregation of probabilities by user-defined cell-level annotation groups, such as cell type annotations, hoodscanR generates a comprehensive probability matrix 𝑃, where each value represents the probability of each cell belonging to a specific cell neighborhood (Figure 1). This matrix describes the cellular neighborhood profiles for all cells, serves as the backbone for downstream analyses, enabling researchers to delve into spatial patterns and relationships.

To investigate how the hyperparameter k and 𝜏 affects the results generated by hoodscanR, we conducted an extensive examination of the probability matrix across a range of k and 𝜏 values (see Methods). This analysis revealed that different k values generate highly similar results, with a mean Pearson correlation coefficient of 0.93 (Supplementary figure S1). Regarding 𝜏, smaller values assign greater weights to nearby cells, while larger 𝜏 values consider more distant cells as contributors to the neighborhood (Supplementary figure S2). Thus, the choice of 𝜏 becomes essentially linked to the specific biological questions being addressed. For example, smaller 𝜏 values, such as one-fifth of the median of the distance matrix, which is set to the default 𝜏 value in the hoodscanR package, are more suitable for analyses focused on local interactions, where nearby cells have stronger influence on the neighborhood calculation. In contrast, larger 𝜏 values, such as the median of the distance matrix, are ideal for capturing more global spatial relationships, incorporating cells that are further away as significant components of the neighborhood.

After neighborhood identification, hoodscanR extends its capabilities to offer a diverse suite of downstream neighborhood analysis tools (Figure 1). Users can apply these tools to visualize spatial relationships, evaluate co-localization patterns, perform spatial neighborhood clustering of cells, and obtain cell-level neighborhood annotations. These functionalities allow researchers to gain insights within the spatial transcriptomic landscape, facilitating the discovery of novel biological knowledge. Last but not least, one of the hallmark features of hoodscanR is using the Bioconductor spatialExperiment infrastructure as the backbone of the analysis. This significantly increases the compatibility of intermediate results from hoodscanR with diverse Bioconductor packages tailored for preprocessing, quality control, normalization, cell type annotation, and various downstream analyses specifically crafted for spatial transcriptomics data. In conclusion, hoodscanR provides a powerful and flexible method for spatial neighborhood identification and analysis.

### Benchmarking hoodscanR in spatial domains identification

Building upon the foundation of cell-level neighborhood probability matrix (Figure 1), hoodscanR allows users to perform unsupervised clustering, grouping cells with similar neighborhood distribution patterns into cohesive clusters. This data-driven approach enables the classification of cells based on their spatial relationships within the tissue slide, identifying neighborhood-driven spatial domains.

To evaluate the effectiveness and robustness of the neighborhood-based spatial domain identification function in hoodscanR, and compared with other tools, we conducted a benchmarking experiment against several state-of-the-art methods (see Methods). This benchmark experiment involved detecting spatial domains of 16 publicly available datasets, covering a range of spatial platforms and tissue types, including CosMx NSCLC (1), MERFISH mouse colon (28), STARmap mouse cortex (29) and Xenium breast cancer (2,30). The datasets were chosen because the tissue is well-annotated with region labels or there are pathological annotataion that can be served as ground truth of spatial domains. hoodscanR was benchmarked against seven other methods that can perform spatial domain detection: BuildNicheAssay from Seurat, Banksy, BayesSpace, MERINGUE, SpaGCN, Stagate and Utag (Figure 2A and supplementary figure S3-7). As a result from these 128 experiments, hoodscanR outperform other methods, followed by Utag and Banksy by achieving the highest performance score on average across all tested datasets (Figure 2B). More importantly, hoodscanR outperforms all others in computing efficiency (Figure 2C), being approximately 21-fold faster on average than Banksy, which ranks second in speed.

**Figure 2.**
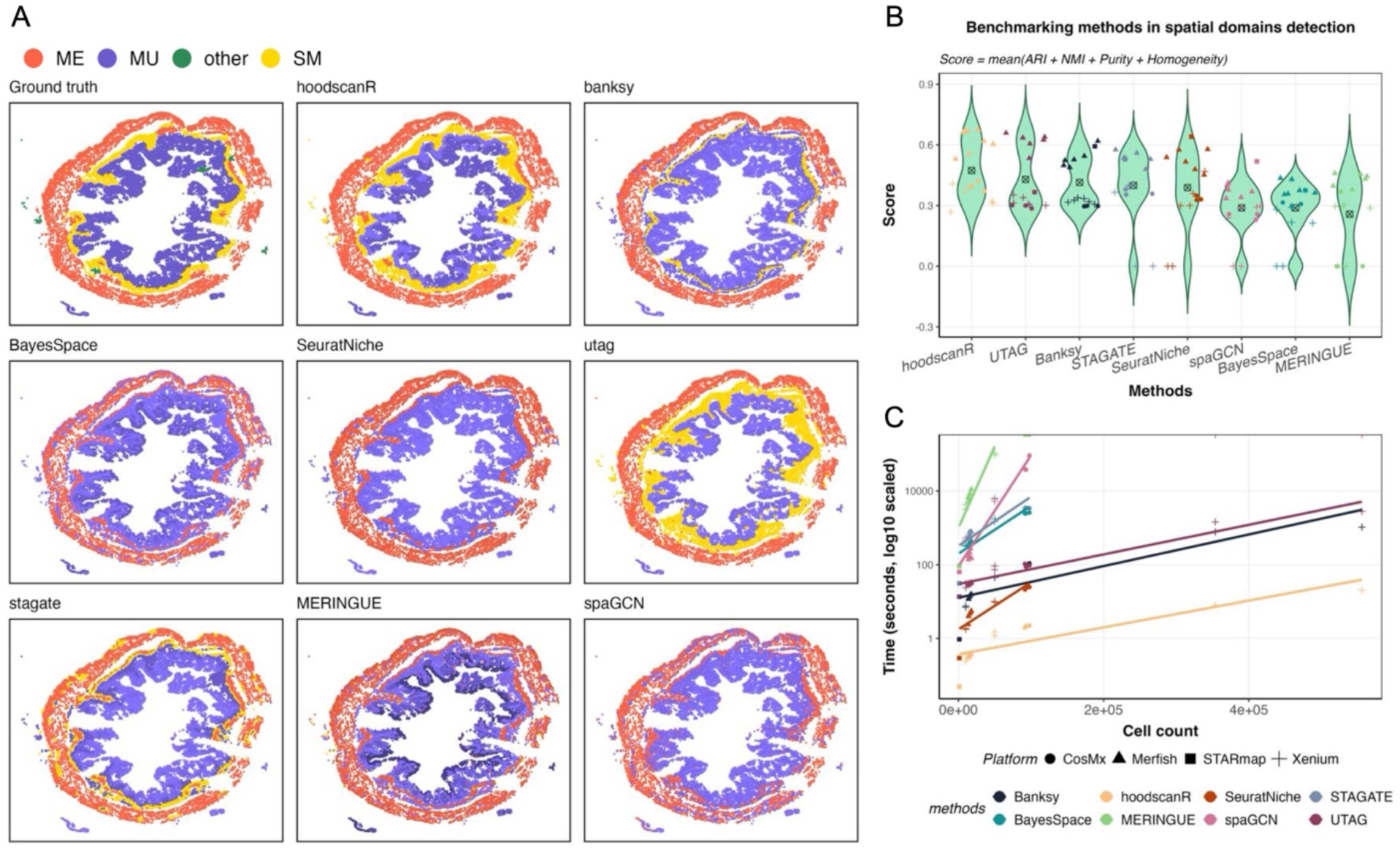
Benchmarking hoodscanR against other methods in detecting spatial domains. A: Spatial maps of the MERFISH mouse colon data, colored by the spatial domains detected from different methods compared to the ground truth domain annotations, including muscularis externa (ME), mucosa (MU), submucosa (SM) and other (top left corner). B: Violin plots showing the performance score of each method across all tested datasets, ordered from the highest to the lowest mean score. C: Computational efficiency of each method, plotted as the log10-scaled time (in seconds) required to process datasets. Shapes represent the platform of the dataset, colors denote the methods, and the lines are generalized linear smooths indicating overall trends for each method.

This advantage in computational speed is particularly important as increasingly large and high-resolution datasets will be generated with the advancements in spatial transcriptomics technologies. Additionally, hoodscanR can recapitulate tissue spatial architecture in a biologically coherent manner. For example, in the MERFISH mouse colon dataset (Figure 2A), hoodscanR accurately delineates four concentric layers, mucosa (MU), submucosa (SM), muscularis externa (ME), and other, all of which contain multiple cell types. This result closely mirrors the ground truth. By clustering cells with similar neighborhood distributions, hoodscanR captures subtle boundaries more effectively than many alternative methods, preserving the colon’s characteristic concentric organisation, such as the ME structure. Taken together, these results highlight strengths of hoodscanR in domain identification across large-scale spatial transcriptomics datasets.

Additionally, to evaluate the robustness of hoodscanR across different resolutions of cell type annotations, we conducted an experiment using high-resolution, medium-resolution, and low-resolution annotations as inputs. The high-resolution annotations included detailed cell types, such as CD4+ T cells, CD8+ T cells, and macrophages. The medium-resolution annotations combined all T cells into a single category, and the low-resolution annotations further grouped all immune cells into a single “Immune” category. Despite the reduction in annotation resolution, the identified neighborhood-based spatial domains have a Normalized Mutual Information (NMI) score of greater than 0.8 when comparing using the high-resolution results as the reference (Supplementary figure S8). Taken together, these results showcase the power of hoodcanR in accurately identifying neighborhood-based spatial domains in a scalable and efficient manner. They also indicate that hoodscanR is robust to variations in annotation granularity, maintaining the integrity of the spatial relationships even when the resolution of cell type annotations is reduced.

### hoodscanR identifies celluar neighborhoods in cancer

To demonstrate the power of hoodscanR in detecting spatial cellular neighborhoods, we performed neighborhood identification on two publicly available spatial transcriptomics datasets obtained from different *in situ* transcriptomic platforms: breast cancer data obtained from the 10X Genomics Xenium (Figure 3A) and non-small cell lung cancer (NSCLC) data acquired from the Nanostring CosMx Spatial Molecular Imager (Figure 3B). We first applied hoodscanR onto the breast cancer dataset using the default parameters (k=100 and 𝜏=median(dist^2)/5). hoodscanR allows us to perform neighborhood identification by profiling neighborhood distributions for each cell within 6 seconds, representing the probability of a cell being situated within each distinct cell-type neighborhood (Figure 3C and 3F).

**Figure 3.**
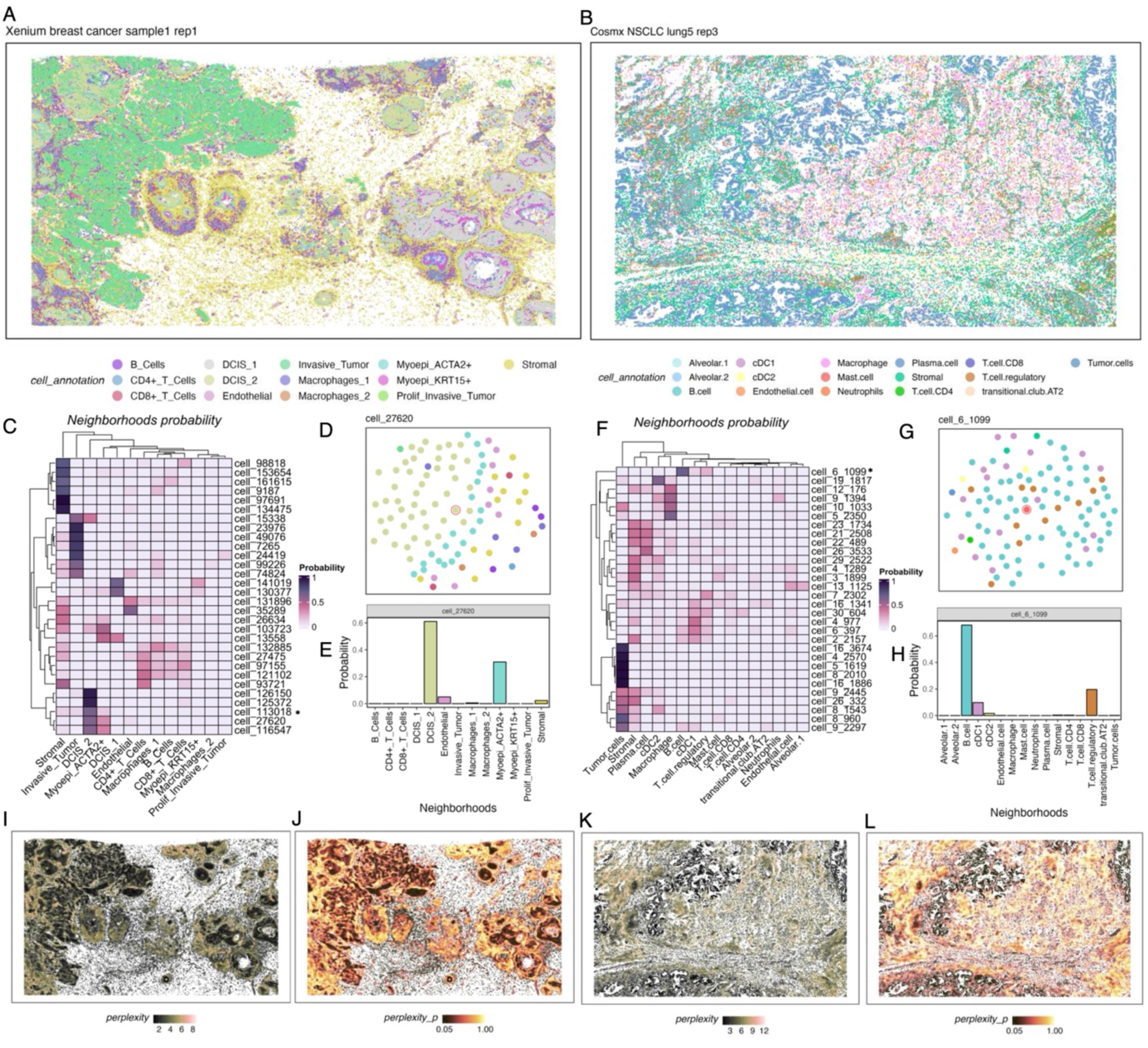
Neighborhood identification in 10X Xenium breast cancer data and Nanostring CosMx NSCLC data. Cell type spatial distribution in the breast cancer data (A) and NSCLC data (B). Neighborhood distribution visualization via heatmap of randomly selected 30 cells from breast cancer data (C) and NSCLC data (F), darker color means higher probability of the cell located in specific cell type neighborhood. The cell type spatial distribution in the spatial area around the selected cells (marked by * in the heatmap) in breast cancer data (D & E) and NSCLC data (G & H). Perplexity spatial distribution of cells in the breast cancer data (I) and NSCLC data (K). P-value distribution of perplexity in the breast cancer data (J) and NSCLC data (L).

To validate the accuracy of hoodscanR in characterizing these cellular neighborhoods, we focus on one randomly selected cell for each dataset by exploring the distribution of cell types within their spatial area (Figure 3D, E, G, and H). For instance, we examined a ductal carcinoma in situ (DCIS) grade 2 cell, an early form of breast cancer cells, from the Xenium data (Figure 3C and D: cell ID 27620), where we observed 47 DCIS grade 2 cells and 21 ACTA2+ myoepithelial cells from the nearest 100 neighboring cells (Figure 3D). Consistently, hoodscanR assigned probabilities of 61.02% for residing in the DCIS grade 2 neighborhood and 30.96% for the ACTA2+ myoepithelial neighborhood for this specific cell (Figure 3E). Similarly, when assessing a stromal cell within the CosMx NSCLC data (Figure 3F: cell ID 6_1099), we observed that hoodscanR assigned probabilities of 68.2% for the B cell neighborhood and 19.67% for the regulatory T cell neighborhood while there are 66 and 11 B cells and regulatory T cells in the nearest 100 neighboring cells (Figure 3G and H). These examples demonstrate the power of hoodscanR in accurately characterizing cellular neighborhoods within spatial transcriptomics data, regardless of the platform, and its capacity to accommodate scenarios where cells may belong to neighborhoods of multiple cell types. The identification of B cell neighborhoods is particularly noteworthy in the context of cancer therapy responses. B cell neighborhoods serve as crucial sites for antibody production, contributing to the immune response against tumor cells and influencing therapeutic efficacy (31). Furthermore, in lung cancer, the presence of tertiary lymphoid structures (TLS), characterized by highly organized T and B lymphocyte colonies within nonlymphoid tissues, has been associated with favorable clinical outcomes in non-small cell lung cancer (NSCLC) (32). These structures, resembling secondary lymphoid organs, play an important role in regulating antitumor immune responses and are emerging as potential targets for novel therapeutic interventions. By delineating cellular neighborhoods, including B-cell-rich TME, hoodscanR offers the potential for investigating the relationship between immune cells and tumor cells within the TME, providing insights that could inform the development of more effective cancer therapies.

Furthermore, hoodscanR introduces an additional analytical dimension by enabling the computation of uncertainty, which is measured by perplexity, and performing permutation test for each cell (see Methods). Perplexity is calculated from the probability matrix, capturing the spatial relationships among cells and their respective neighborhoods. Perplexity provides overall measurement of the uncertainty and complexity of cell neighborhoods (Figure 3I and K). This in turn reveals regions of the TME with distinct cellular compositions and areas with complex interactions between cell types. P-values of perplexity (Figure 3J and L) can be obtained via an empirical permutation test (see Methods). This allows users to identify regions with significant higher perplexity from tissue. Altogether, these metrics provide an understanding of the heterogeneity and complexity present within tissues, allowing researchers to gain novel insights and make discoveries in the spatial transcriptomics landscape.

### hoodscanR performs neighborhood-based downstream analyses

Existing neighborhood identification methods, such as Squidpy and Giotto, predominantly focus on neighborhood co-localization analyses. Another key function of hoodscanR is to generate neighborhood profile at single-cell level and to carry out neighborhood-based downstream analyses, features notably absent in other existing tools. To demonstrate the versatility of hoodscanR, we use the CosMx NSCLC dataset as an example.

Firstly, as with other spatial analysis tools, hoodscanR can perform neighborhood co-localization analysis by computing Pearson correlations on the neighborhood distribution of cells. Hence, the co-localization status of each cell type neighborhood within this specific tissue slide can then be visualized (Figure 4A). To benchmark the ability of hoodscanR, Squidpy and Giotto when carrying out co-localization analysis, we subset the Xenium breast cancer data and CosMx NSCLC into different resolution, followed by applying these tools to the subsets. As a result, hoodscanR demonstrated superior computational performance (Supplementary figure S9), while delivering similar outcome (mean Pearson correlation coefficient of 0.781) of neighborhood co-localization compared to both Xenium and CosMx data (Supplementary figure S10 and S11). The computational efficiency gains significance, particularly in the context of the growing spatial data resolutions and larger tissue areas.

**Figure 4.**
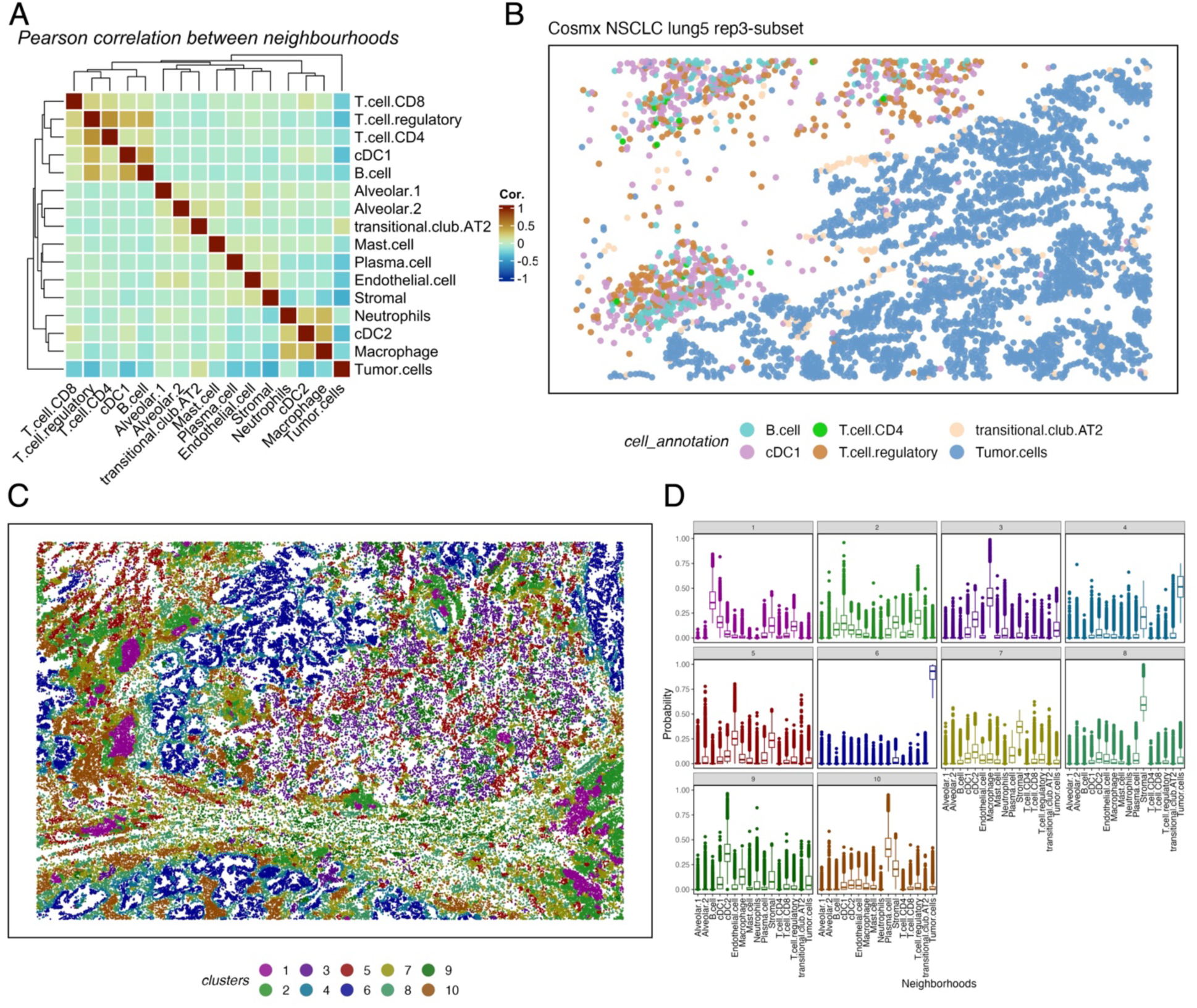
Neighborhood-based downstream analyses in CosMx NSCLC data performed by hoodscanR. A: heatmap representing the co-localization status of cell type neighborhood in this tissue slide. Colors denote positive (dark red) and negative (blue) correlations. B: Spatial location plot of a subset of cell types in this slide. Colors denote cell types. C: Spatial location plot of the slide with colors stratified by neighborhood-based clusters. D: The neighborhood distribution profiles of each neighborhood-based cluster identified in the tissue slide.

To validate the co-localization results, we present a subset of cell types alongside their spatial distribution within the breast cancer tissue slide, showing that immune cell types such as B cells, T cells, and macrophages exhibit co-localization, while they are distinctly separated from tumor cells (Figure 4B). This observation serves as robust validation of the co-localization analysis results generated by hoodscanR, thus reinforcing the effectiveness and reliability of hoodscanR in revealing spatial relationships within various tissue environments, particularly when dealing with complex spatial transcriptomics data.

In the CosMx NSCLC data, we applied unsupervised clustering to delineate 10 distinct clusters (see Methods), each representing a unique spatial pattern within the tissue (Figure 4C), demonstrating complex spatial associations. For instance, cluster 1, a candidate cluster for TLS, corresponds to a neighborhood including B cells, cDC1 cells, and stromal cells (Figure 4D - 1), cluster 3 aligns with macrophages and cDC2 cells (Figure 4D - 3), and cluster 6 corresponds to tumor cells (Figure 4D - 6). This unsupervised clustering approach facilitates the identification of diverse cellular neighborhoods and their unique spatial signatures, providing a comprehensive view of the complexity of TME.

### hoodscanR detects changes between tumor cells from different neighborhoods

Building on these findings, we then used uniform manifold approximation and projection (UMAP) to perform dimension reduction visualization on the expression data of the CosMx NSCLC data, enabling the projection of gene expression profiles into a lower-dimensional space. This facilitates the visualization of cell lineage (Figure 5A) alongside identified neighborhood-based clusters (Figure 5B). A distinctive feature observed is the dispersion of cells of the same cell type across different neighborhood clusters, signifying diverse spatial neighborhood profiles. For example, a substantial proportion (76.14%) of macrophages (pink points in Figure 5A) are distributed across various neighborhood clusters, including cluster 3 (43.63%), indicative of the macrophage + cDC2 + tumor neighborhood, clusters 7 (12.31%), and cluster 9 (20.2%), representing the stromal + cDC2 neighborhood (Figure 3D and 5B).

**Figure 5.**
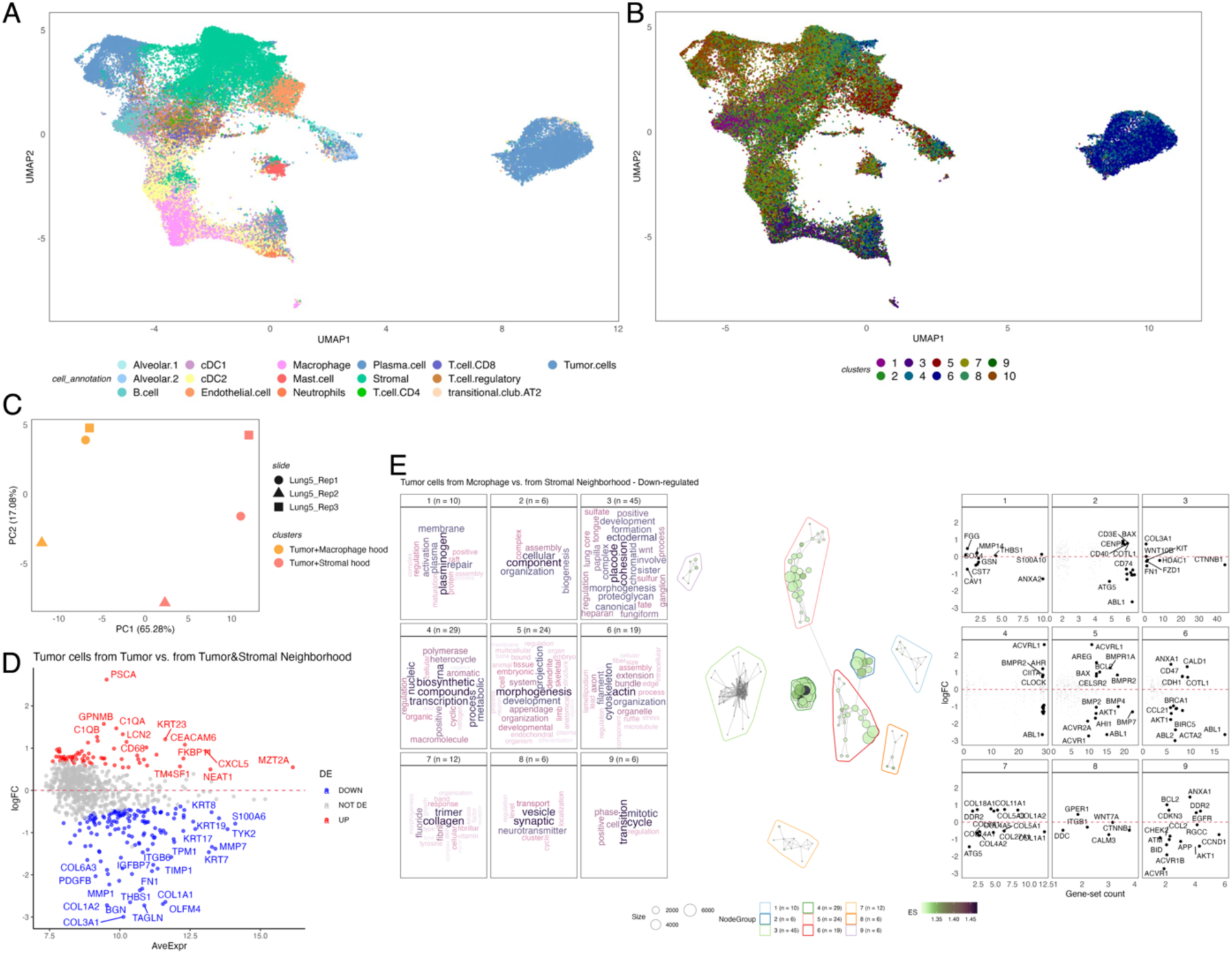
Neighborhood-based transcriptomics analysis in CosMx NSCLC data. A: UMAP of expression of cells in the data, colors denote cell types. B: UMAP of expression of cells in the data, colors denote the identified neighborhood-based clusters. C: PCA of pseudo-bulk samples of tumor cells from two different neighborhood clusters across three consecutive slides. Colors denote clusters and shapes denote replicates. D: MA plot describing the outcome of the DE analysis. Colors indicate up- (red) or down-regulated (blue) genes. E: vissE visualization of significantly enriched gene-sets from the down-regulated DE genes in the comparison between tumor cells from macrophage neighborhood and tumor cells from stromal neighborhood. Left panel are word cloud plots describing gene-set clusters of different biological themes, middle panel is the gene-set overlap network graph of gene-sets, and right panel is the fold-change (log2-scaled) for genes belonging to each gene-set cluster.

An important aspect of hoodscanR lies in its ability to investigate the relationship between spatial neighborhoods and transcriptional changes. To demonstrate this, we conducted a nuanced analysis by extracting and pseudo-bulking tumor cells from two distinct neighborhood clusters: the stromal cluster and the macrophage cluster across three consecutive slides (Supplementary figure S12). Interestingly, different spatial neighborhoods contribute significantly to the variation observed in the first dimension from a principal component analysis (PCA) of expression of pseudo-bulk samples (Figure 5C). This signifies that hoodscanR can capture transcriptional changes attributed to diverse spatial neighborhoods.

Subsequently, we performed a differential expression (DE) analysis using the limma-voom pipeline (24), identifying 220 DE genes from 832 genes, including 73 up-regulated and 147 down-regulated genes when comparing tumor cells from the macrophage neighborhood to those from the stromal neighborhood (Figure 5D and supplementary file 1). Additionally, we performed gene-set enrichment analysis on the identified DE genes with MSigDB gene-sets (33,34), detecting 384 significantly enriched gene-sets (Supplementary file 2). We further perform unsupervised clustering on gene-sets using vissE (26), identifying clusters of gene-sets networks (Figure 5E and supplementary figure S13, middle panel). Notably, we observe pathways enriched in down-regulated DE genes related to collagen, such as *collagen-activated tyrosine kinase receptor signalling pathway* and *collagen metabolic and catabolic process* (Figure 5E left panel). These pathways are accompanied by the differential expression of key collagen-related genes, such as *COL1A1*, *COL11A1* and *COL5A1* (Figure 5E right panel). Previous studies have found that the overexpression of collagen genes such as *COL11A1* (35) and *COL3A1* (36) in NSCLC may indicate poor prognosis and drug resistance, and *COL1A1* is correlated with immune infiltration in NSCLC (37). Our finding of these genes that are expressed significantly more in the tumor cells from the macrophage neighborhood than tumor cells from the stromal neighborhood can potentially lead a more detailed investigation of the biological mechanism about transcriptional changes within the context of cancer spatial TME studies. In essence, hoodscanR introduces a novel perspective in spatial transcriptomics, allowing the identification of transcriptomic changes across subtly different spatial neighborhoods and providing insights into the spatial organisation within the TME.

### hoodscanR supports different gene annotations

The accuracy of cell type-based cellular neighborhoods, as identified in the previous case, is inherently tied to the accuracy of cell type annotation. Thus, we built hoodscanR to be flexible and capable to detect various cellular neighborhoods based on different gene annotation inputs. A particularly valuable application is gene expression-based neighborhoods detection. To showcase this, we utilised a breast cancer tissue, where we can focus on breast cancer-related hormone receptor genes, including androgen receptor gene (*AR*), estrogen receptor gene (*ESR1*), and progesterone receptor gene (*PGR*). By assessing if these genes are expressed or not, we classified 574,527 cells from a Xenium Invasive ductal carcinoma (IDC) dataset into eight distinct groups (Figure 6A). Similar to the previous neighborhood identification based on cell types, hoodscanR can identify spatial domains based on gene expression-specific neighborhoods (Figure 6B and supplementary figure S14). These analyses lead to a nuanced understanding which adeptly discover tumor cells located within neighborhoods characterized by varying combinations of hormone receptors. This not only suggests a spatial perspective on the progression of DCIS, influenced by distinct combinations of hormone receptors but also sheds light on the higher-order spatial structure of cells with different hormone receptor expression profiles.

**Figure 6.**
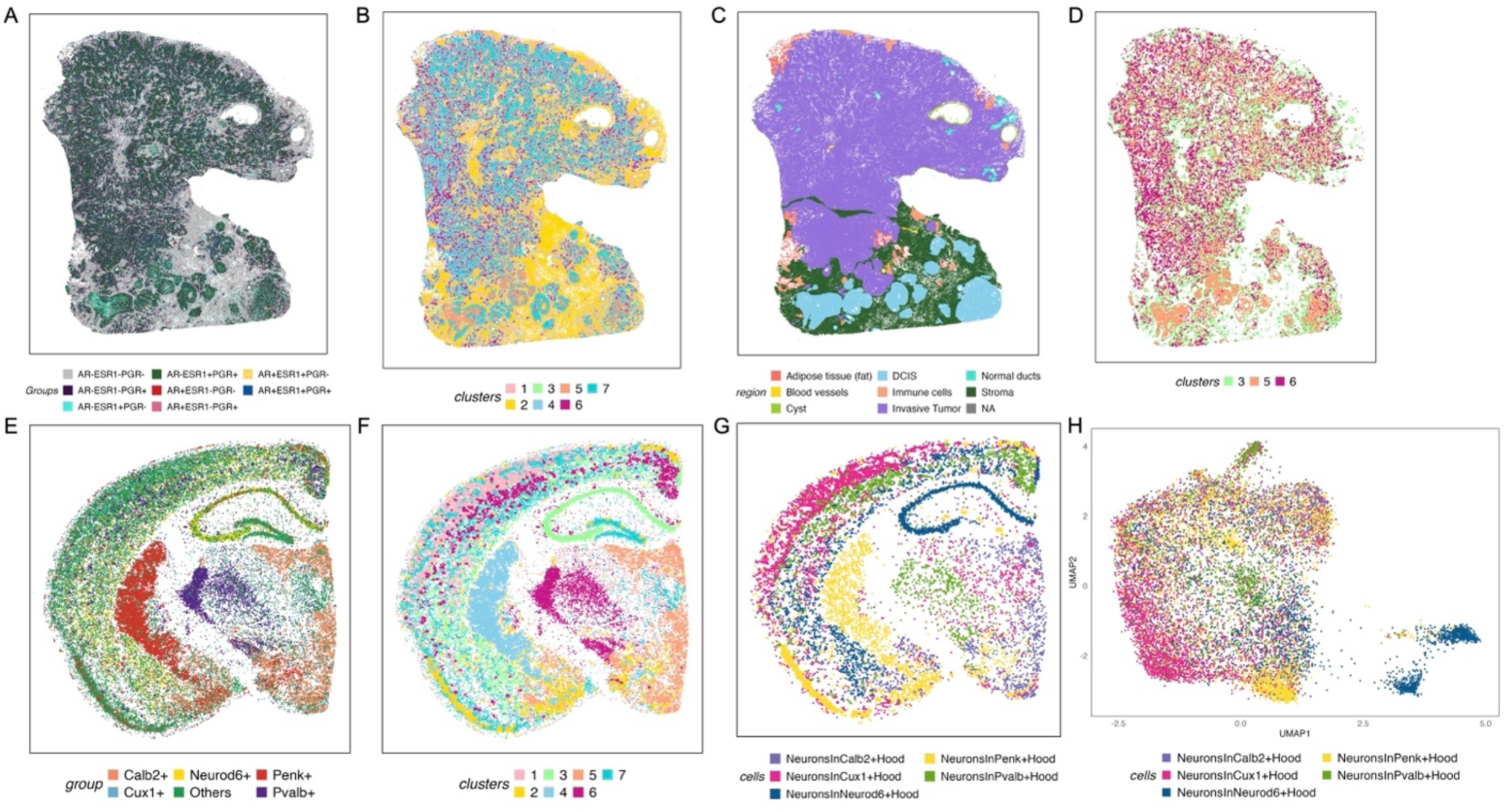
Gene expression-based neighborhood analysis in 10X Xenium IDC (A-D) and VisiumHD mouse brain (E-H) data using hoodscanR. A: spatial plot showing the hormone gene expression-based cell grouping on the tissue slide. B: gene expression-based domains in IDC tissue. C: pathological annotation of regions on the tissue. D: spatial plot showing specific neighborhood clusters, including cluster 3, 5 and 6. E: spatial plot showing the selected gene expression-based cell grouping. F: gene expression-based domains in mouse brain tissue. G: neurons spatial distribution in different gene-expression neighborhoods. H: a dimension reduction UMAP visualization of the neurons from different neighborhoods.

Moreover, interesting insights can be observed by comparing the pathological annotations (Figure 6C) of this slide (30), where invasive and non-invasive (DCIS) cancer phenotype regions were annotated, with the identified neighborhood clusters. Invasive tumors exhibit a distinctive pattern with cells from cluster 3 scattered throughout, accompanied by cells from cluster 6 (Figure 6D). Both clusters are associated with *ESR1*+*PGR*+ and *ESR1*-*PGR*+ cell neighborhoods, respectively. This finding aligns with previous findings indicating that nearly 80% of invasive breast cancers are ER-positive, and PR is overexpressed in ER+ tumors (38). Conversely, DCIS regions are notably associated with cells from cluster 5, which form the inner layer surrounded by cells from cluster 3, comprising the outer layer (Figure 6D). Cluster 5 includes cells expressing *AR*, which is consistent with previous findings showing *AR* expression in DCIS components adjacent to invasive cancer (39). Additionally, the expression of *AR* has been reported to decrease as the disease progresses from DCIS to invasive cancer (40). These observations suggest a potential relationship between the tumor type (non-invasive or invasive) and the higher-order spatial organisation of cells with diverse hormone receptor expression profiles.

To further demonstrate the broad application of hoodscanR beyond cancer and human data, we analysed the publicly available Visium HD mouse brain data. In this case, we selected five marker genes that can divide the brain into different regions: *Calb2* for paraventricular nucleus of the thalamus (PVT) (41), *Neurod6* for deeper layers of the cortex (42), *Penk* for striatal medium spiny neuron (43), *Cux1* for upper layer of the cortex (44) and *Pvalb* for hippocampus (45). Additionally, we included the gene *Rbfox3*, which is exclusively expressed in neuron cells (46). The spatial map of these cells reveals their distinct regional expression patterns within the brain tissue (Figure 6E and supplementary figure S15). Applying hoodscanR, we can identify neighborhood-based clusters (Figure 6F and supplementary figure S16). These clusters largely correspond to specific brain regions (Supplementary figure S17), reflecting the well-structured nature of the brain tissue, with each cluster predominately associated with one of the maker genes.

Focusing on neuron cells by filtering the dataset for cells expression *Rbfox3*, we explored how these neurons are distributed across different neighborhoods (Figure 6G). By conducting dimension reduction via UMAP on the expression data of these neurons (Figure 6H), we can visualise the neurons from different neighborhoods tend to cluster differently, indicating variance in expression between these neurons, especially between those within Neurod6+ neighborhoods and Calb2+ neighborhoods (Supplementary figure S18). Moreover, performing a differential expression analysis between neurons from these two neighborhoods revealed a set of genes that are significantly upregulated in the Neurod6+ region compared to the Calb2+ region (Supplementary figure S19), indicating distinct microenvironmental influences and potential functional specialization. For example, *Nrgn*, *Hpca*, and *Rasgrp1*, which are all involved in regulating intracellular calcium signalling and synaptic plasticity, are up-regulated in the neurons from the Neurod6+ neighborhood, predominantly localized to the hippocampal area. This observation aligns with previous studies showing that these genes are critical for synaptic plasticity (47–50), which underlies learning and memory. Taken together, these patterns illustrate how local spatial neighborhood composition can shape neuronal identity and function, highlighting hoodscanR’s ability to detect spatially restricted transcriptional differences even in highly structured tissues like the brain.

In summary, these findings highlight hoodscanR’s capability to identify subtle spatial changes in expression-based cellular neighborhoods, providing novel insights into the complex spatial dynamics of gene expression in both cancerous and non-cancerous tissues.

## Discussion

Spatial technologies are pushing the limits toward profiling spatial transcriptomics at single-cell level. To make the best use of these cutting-edge technologies, we developed hoodscanR, a powerful yet user-friendly Bioconductor package for revealing spatial cellular relationships within high-dimensional spatial transcriptomics datasets via spatial cellular neighborhoods identification and neighborhood-based downstream analyses. In our benchmarking experiments, hoodscanR demonstrated robust accuracy and computation efficiency compared to several state-of-the-art methods, further validating its utility in diverse spatial transcriptomics applications.

While we have shown hoodscanR can identify biologically meaningful cellular neighborhoods, as with all methods it is not without limitations. Firstly, although hoodscanR is flexible in relation to gene annotations, the preprocessing of spatial transcriptomics data is essential. Current quality control procedures of spatial transcriptomics data predominantly operate at cell level, potentially missing crucial details detectable only at a transcript (subcellular) level. A more comprehensive strategy for data preprocessing, accounting for information at transcript level, filtering out uninformative cells accurately and enhancing the precision of neighborhood identification, becomes imperative. Secondly, hoodscanR is a cell-based method, emphasizing the critical role of accurate cell segmentation, which hoodscanR depends upon other methods. The segmentation process, which determines how individual cells are identified and their spatial coordinates are established, is fundamental to the accurate detection of cellular neighborhoods. Variations in segmentation methods, such as differences in how cell boundaries are defined or how centroids are calculated, can lead to significant differences in cell type distributions and spatial relationships within the tissue. Such variations can impact the neighborhood detection results produced by hoodscanR, potentially leading to different biological interpretations. A systematic review of existing segmentation methods is lacking, necessitating future research to evaluate and compare methods under diverse spatial transcriptomic platforms. Lastly, while hoodscanR enables exploration of spatial gene expression patterns, users must interpret results cautiously. Recognizing that spatial context alone may not capture the full complexity of molecular interactions within a tissue. Integrating multi-omics data can provide a more comprehensive understanding, ensuring spatial analyses are embedded within a broad molecular context. Strategically addressing these considerations allows researchers to fine-tune the utilization of hoodscanR, strengthening the integrity of their analyses and facilitating the discovery of novel insights.

Three of the major strengths of hoodscanR are its compatibility, adaptability and flexibility. Developed based on the Bioconductor SpatialExperiment infrastructure, hoodscanR exhibits compatibility with many other spatial and single-cell RNA-seq-based tools from Bioconductor, amplifying its utility for downstream analyses.

Importantly, its adaptability makes it platform agnostic, demonstrated by successful applications on Nanostring CosMx, 10X Genomics Xenium, MERFISH and STARmap. Spatial datasets from various platforms, including Vizgen MERSCOPE, 10X Visium HD, BGI STOmics, and Akoya Biosciences CODEX, can also undergo comprehensive analysis using hoodscanR, given the availability of cell-based coordinates. The flexibility of hoodscanR is demonstrated by the types of annotations it can use. With the Xenium breast cancer data, hoodscanR showcased this by not only using cell type annotations but also accommodating gene expression level grouping of cells, suggesting the potential for exploring additional annotation options such as ligand-receptor or growth factor-receptor annotations. This design provides researchers the flexibility to generate hypotheses on the basis of integrating spatial localization and gene expression before testing statistical associations between these. Lastly, while hoodscanR is not designed for direct cell-cell communication analysis, it plays a crucial role in accurately identifying and characterizing spatial neighborhoods. This capability can complement existing cell-cell communication tools, such as COMMOT (51), CellChat (52) or CellPhoneDB (53), by providing a more refined spatial context that may enhance the accuracy and robustness of cell-cell communication network identification in complex tissue environments.

The significance of hoodscanR in identifying and analyzing neighborhoods becomes particularly crucial in the context of complex diseases, notably in cancer research. Neighborhood information is indispensable for unraveling complex disease etiology, especially so for understanding disease progression. In cancer research, where the TME plays an important role in dictating therapy responses, the ability offered by hoodscanR to identify neighborhoods offers a unique perspective to investigate novel mechanisms underlying the transition of cancer cells at both transcriptomic and proteomic level. Notably, our findings from the Nanostring CosMx NSCLC dataset identified cellular neighborhoods that are potentially associated with the presence of TLS, which correlated with positive clinical results (32). This capability offers the potential to contribute valuable insights into the progression of cancer, paving the way for the development of novel therapeutic strategies.

In conclusion, our study introduces hoodscanR, a Bioconductor package designed for comprehensive neighborhood analysis in spatial transcriptomics datasets. Through its integration with the SpatialExperiment infrastructure and efficient algorithms, hoodscanR offers fine-tuned control over neighborhood identification, allowing researchers to investigate complex cellular relationships within spatially resolved datasets. By demonstrating hoodscanR’s efficacy on the 10X Genomics Xenium breast cancer and Nanostring Technologies CosMx non-small cell lung cancer datasets, we showcase its ability in identifying cellular neighborhoods and elucidating spatial gene expression patterns. Furthermore, our findings emphasize the significance of neighborhood analysis in understanding the complex TME of cancer tissues, which can potentially lead to the identification of novel biological mechanisms underlying disease progression and therapeutic responses. Importantly, hoodscanR’s flexibility in handling diverse spatial datasets and its ability to accommodate different types of cell annotations enhance its utility for a wide range of spatial transcriptomic studies. Overall, hoodscanR contributes to advancing the field of spatial transcriptomics by providing researchers with a powerful tool, thereby paving the way for deeper insights into tissue biology and disease mechanisms.

## Data availability

The Nanostring CosMx non-small cell lung cancer (NSCLC) data utilized in this study was sourced from the official Nanostring website: https://nanostring.com/products/cosmx-spatial-molecular-imager/ffpe-dataset/nsclc-ffpe-dataset/. The CosMx dataset was generated from FFPE NSCLC tissue samples using a 960-plex CosMx RNA panel. The 10X Xenium breast cancer datasets used in this study were retrieved from the 10X publicly available database at https://www.10xgenomics.com/datasets. The Xenium *in situ* dataset comprises human breast cancer FFPE sections and utilizes a 280-gene Xenium Human Breast Gene Expression Panel supplemented with 33 additional custom genes. The MERFISH mouse colon data was downloaded from https://doi.org/10.5061/dryad.rjdfn2zh3, originated from Cadinu et al., 2024 (28). The STARmap mouse cortex data was sourced from the Wang et al., 2018 study (29). The 10X VisiumHD mouse brain dataset was sourced from the official 10X Genomics website: https://www.10xgenomics.com/datasets/visium-hd-cytassist-gene-expression-libraries-of-mouse-brain-he.

## Code availability

The code used for performing the described analyses is available in GitHub at https://github.com/ningbioinfo/hoodscanR_manuscript_code. The hoodscanR package is freely available in Bioconductor (release > 3.18) at https://bioconductor.org/packages/release/bioc/html/hoodscanR.html.

## Acknowledgements

The authors thank both Nanostring Technologies and 10X Genomics for releasing publicly available CosMx and Xenium datasets, respectively.

## Author information

Conceptualization: N.L., J.M., C.W.T. and M.J.D. Method development: N.L., D.D.B., A.M., M.L. Writing – original draft: N.L. Writing – draft revision: N.L., J.C., S.C.L., M.K., J.C., A.K., Y.C., C.W.T., F.L, J.M.P.

## Ethic declarations

Not Applicable.

